# ClinGen Expert Clinical Validity Curation of 164 Hearing Loss Gene-Disease Pairs

**DOI:** 10.1101/534040

**Authors:** Marina T. DiStefano, Sarah E. Hemphill, Andrea M. Oza, Rebecca K. Siegert, Andrew R. Grant, Madeline Y. Hughes, Brandon J. Cushman, Hela Azaiez, Kevin T. Booth, Alex Chapin, Hatice Duzkale, Tatsuo Matsunaga, Jun Shen, Wenying Zhang, Margaret Kenna, Lisa A. Schimmenti, Mustafa Tekin, Heidi L. Rehm, Ahmad N. Abou Tayoun, Sami S. Amr, on behalf of the ClinGen Hearing Loss Clinical Domain Working Group

**Author notes:** These authors contributed equally to this effort (Marina T. DiStefano, Sarah E. Hemphill). Send all manuscript correspondence to: Sami Amr, 1-617-768-8377.

## Abstract

**Purpose:** Proper interpretation of genomic variants is critical to successful medical decision making based on genetic testing results. A fundamental prerequisite to accurate variant interpretation is the clear understanding of the clinical validity of gene-disease relationships. The Clinical Genome Resource (ClinGen) has developed a semi-quantitative framework to assign clinical validity to gene-disease relationships.

**Methods:** The ClinGen Hearing Loss Gene Curation Expert Panel (HL GCEP) uses this framework to perform evidence-based curations of genes present on testing panels from 17 clinical laboratories in the Genetic Testing Registry. The HL GCEP curated and reviewed 142 genes and 164 gene-disease pairs, including 105 nonsyndromic and 59 syndromic forms of hearing loss.

**Results:** The final outcome included 82 Definitive (50%), 12 Strong (7%), 25 Moderate (15%), 32 Limited (20%), 10 Disputed (6%), and 3 Refuted (2%) classifications. The summary of each curation is date stamped with the HL GCEP approval, is live, and will be kept up-to-date on the ClinGen website (https://search.clinicalgenome.org/kb/gene-validity).

**Conclusion:** This gene curation approach serves to optimize the clinical sensitivity of genetic testing while reducing the rate of uncertain or ambiguous test results caused by the interrogation of genes with insufficient evidence of a disease link.

## Introduction

Accurate interpretation of genomic variants is critical for diagnostic utility. According to OMIM, approximately 1738 gene-disease relationships were discovered between 2010 and 2016.^1^ Variants in a gene cannot be clinically interpreted if a gene has not been previously implicated in disease.^2^ Thus, variant interpretation relies on an understanding of the clinical validity of the affected gene. The Clinical Genome Resource (ClinGen),^3^ an NIH-funded initiative building an authoritative central resource to define the clinical relevance of genes and variants for use in precision medicine and research, has developed a semi-quantitative framework to assign clinical validity to gene-disease relationships.^4^ This framework involves the curation of primary published literature to score genetic and experimental evidence, which supports the assignment of a clinical validity classification (Definitive, Strong, Moderate, Limited, Disputed, Refuted, or No Evidence). Conditions known to have a high degree of genetic heterogeneity, such as hearing loss, have hundreds of genes reported as causal in the literature and stand to benefit from this framework to disambiguate gene involvement in disease.

Hearing loss affects approximately 2-3 out of 1000 infants and half of these cases have a genetic etiology.^5^ The auditory system is highly complex, and genetic hearing loss is highly heterogeneous.^6^ There are over 100 genes proposed to be associated with nonsyndromic hearing loss (NSHL) and over 400 genes associated with syndromic forms of hearing loss (Hereditary hearing loss homepage; http://hereditaryhearingloss.org).^7^ Therefore, transparent and systematic evaluations of gene-disease relationships are required for genetic testing to identify the basis of hearing loss in affected individuals or families. Towards this goal, ClinGen assembled a group of experts to form the ClinGen Hearing Loss Clinical Domain Working Group (CDWG) in June of 2016 (http://tinyurl.com/ClinGenHearingLoss).^8^ Along with specifying the ACMG/AMP guidelines for interpretation of variants in hearing loss genes under the Hearing Loss Variant Curation Expert Panel (HL VCEP)^8^ one of the goals of this working group is to assess the clinical validity of genes associated with hearing loss using the ClinGen gene curation framework. The Hearing Loss Gene Curation Expert Panel (HL GCEP) therefore conducted expert curation and review of the clinical validity of 142 genes with 164 total gene-disease relationships, including 105 with NSHL and 59 with syndromic forms of hearing loss. These expert-reviewed curations are publicly available on the ClinGen website (www.clinicalgenome.org).

## Materials and Methods

### Generating a gene list

In total, 142 genes were curated by the working group (Supplementary table 1). This gene list was constructed by aggregating the genes on next-generation sequencing panels for hearing loss from 17 international and US-based laboratories (ARUP, Asper (Estonia), Blueprint (Finland), CeGaT (Germany), Centogene (Germany), CGC Genetics (Portugal), Children’s Hospital of Philadelphia, Cincinnati Children’s, Emory Genetics Laboratory, Fulgent, Genetaq, Greenwood Genetics, Knight, Molecular Otolaryngology and Renal Research Laboratories (MORL), Otogenetics, Partners Healthcare Laboratory for Molecular Medicine, and Prevention Genetics) in the Genetic Testing Registry (GTR; https://www.ncbi.nlm.nih.gov/gtr/, accessed Feb 1 2018). The hearing loss panels from these laboratories were each comprised of at least 20 genes. When a laboratory had multiple panels and no single panel was comprehensive, the gene lists of multiple panels were combined. The number of times each gene appeared on a panel was recorded (Supplementary Table 1).

Five additional genes, *KIAA1199*, *DMXL2*, *TMTC2*, *TUBB4B*, and *SLC44A4*, were not present on any hearing loss panels in the GTR but were included by recommendation of CDWG members.

### Pre-curation and lumping and splitting

OMIM and PubMed were used to search for asserted disease relationships for each gene. If a gene was associated with more than one disease, the diseases were either lumped together or split and curated separately. All genes with a published relationship with NSHL were curated fully with respect to that phenotype. Genes associated with both autosomal dominant (AD) and autosomal recessive (AR) hearing loss were curated separately with respect to each mode of inheritance. Genes linked with one or more hearing loss syndromes underwent pre-curation to identify all possible associated diseases which were reviewed by members of the HL GCEP with clinical expertise. Pre-curation involved a literature search to collect the following information for each gene-disease relationship: 1) If hearing loss is a diagnostic feature of the syndrome, 2) if hearing loss is ever the presenting feature of the syndrome, 3) the penetrance of hearing loss in individuals with pathogenic variants in the gene, 4) the age of onset of hearing loss, 5) the severity, progression, and audiogram shape of the hearing loss (when available) and 6) if individuals with isolated hearing loss were evaluated to rule out the presence of other features of the syndrome (Supplementary Table 2). Syndromic hearing loss conditions only underwent full primary curation if hearing loss had ever been the presenting feature of the syndrome or the additional features could be overlooked during clinical evaluation. For example, the gene *DIAPH1* is linked to AD hearing loss with macrothrombocytopenia, a blood phenotype which can be overlooked during clinical evaluation. For genes linked with multiple hearing loss syndromes, or both syndromic and nonsyndromic hearing loss, curations were either lumped or split per the ClinGen Lumping and Splitting guidelines (https://www.clinicalgenome.org/site/assets/files/9703/lumping_and_splitting_guidelines_gene_curation_final.pdf). If associations with any individual phenotypes within a syndrome had been Disputed or Refuted, they were split from the primary disease relationship in order to highlight the conflicting evidence. Examples of this process are provided in the results.

### Curation and Expert Review

Once the gene list and disease relationships were determined, each gene-disease relationship underwent primary curation by a single curator, using the ClinGen framework as described {https://www.clinicalgenome.org/site/assets/files/8891/gene_curation_sop_2016_version_5_11_6_17.pdf}.^4^

A dual review process was initially used to standardize application of the ClinGen framework: Following primary curation, a secondary curator with expertise in the hearing loss field would review the curation and recommend changes to scoring. The curation was then presented to the full working group. Following these presentations, point assignments and overall classifications were modified when appropriate based on input from the ClinGen HL GCEP. After the first 30 curations, the process became standardized and the secondary review was eliminated, with all curations directly presented to the full committee for review. For well-established gene-disease relationships with an overwhelming amount of evidence, a streamlined review process was used in which curation results were reviewed by one chair of the HL GCEP (Abou-Tayoun, Amr, or Rehm).

Upon expert approval, the curations were approved and published to the ClinGen website (Figure 1, Supplementary Table 1) (https://search.clinicalgenome.org/kb/gene-validity). Detailed scoring modifications to the ClinGen framework made by the HL GCEP, such as downgrading missense variants in the case of consanguinity, are listed in Supplementary Table 3. The two mitochondrial gene curations are not available online, but are included in the supplement (Supplementary Figures 1 and 2).

**Figure 1:**
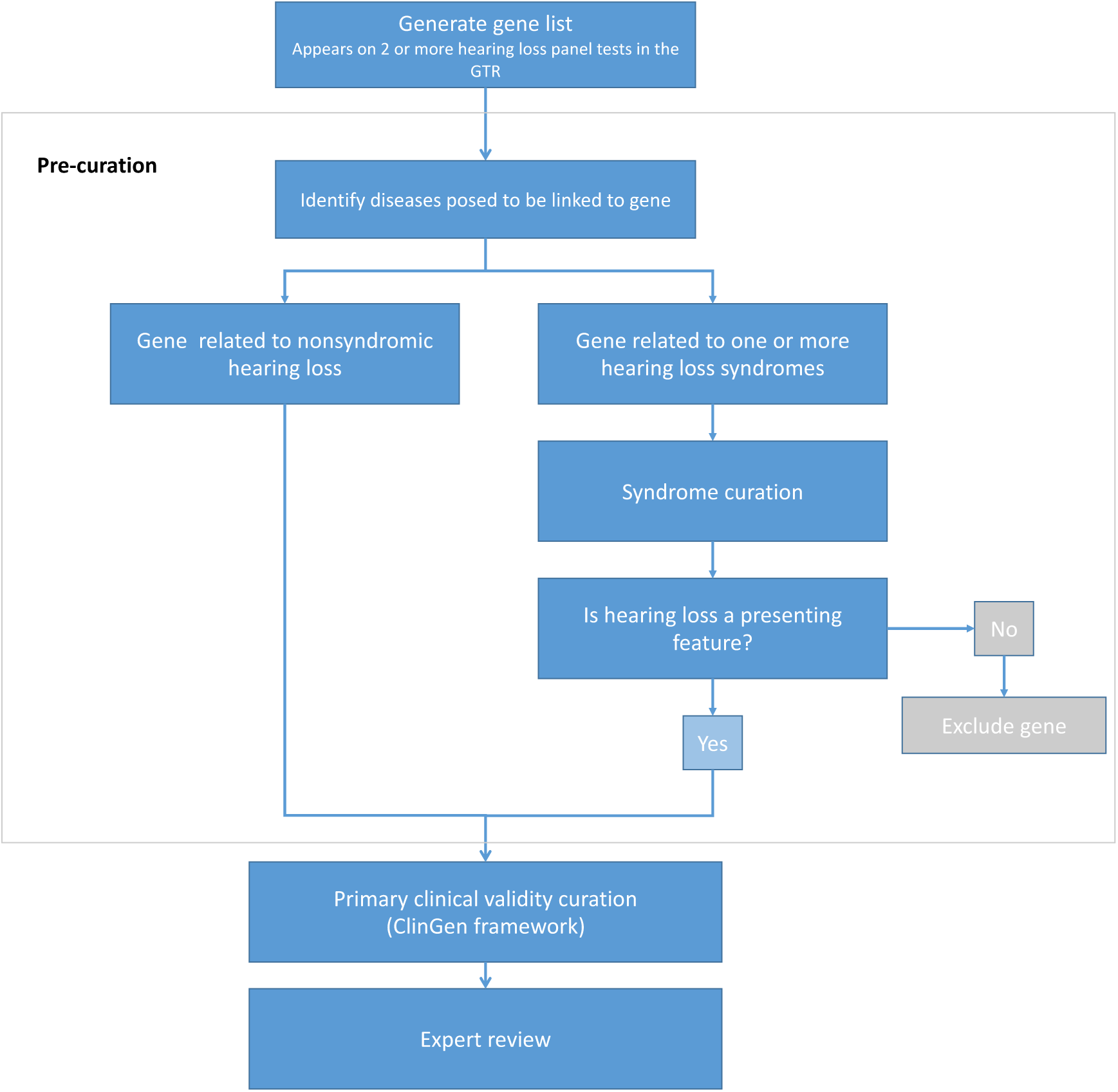
Gene curation workflow. A gene list was generated from 17 clinical testing labs present in the GTR. Nonsyndromic and syndromic genes with hearing loss as a presenting feature were prioritized and fully curated. Syndromic conditions were partially curated in Supplementary Table 2.

## Results

### Curation gene list

The 142 genes analyzed were reported as causal for a broad range of nonsyndromic and syndromic manifestations that are characteristic of the phenotypic heterogeneity and variable expressivity associated with hearing loss. A number of these genes (n=19) had more than one disease claim based on phenotype (nonsyndromic vs. syndromic) or inheritance pattern (AR vs. AD), and each of these claims were reviewed and evaluated separately. An overview of the inheritance patterns reported for each nonsyndromic versus syndromic gene-disease pair is provided in Table 1.

**Table 1.** Condition type (i.e. syndromic vs. nonsyndromic) by inheritance pattern. Curations were performed separately for genes with sufficient evidence to split by condition/inheritance pattern. Counts are representative of gene-disease pairs.

| Inheritance pattern | Nonsyndromic | Syndromic |
| --- | --- | --- |
| Autosomal recessive | 62 | 34 |
| Autosomal dominant | 38 | 21 |
| X-linked | 3 | 4 |
| Mitochondrial | 2 | 0 |

### HL GCEP clinical validity classifications

The ClinGen HL GCEP curated 142 genes and 164 gene-disease pairs, which resulted in 82 Definitive (50%), 12 Strong (7%), 25 Moderate (15%), 32 Limited (20%), 10 Disputed (6%), and 3 Refuted (2%) classifications (Figure 2A, Supplementary Table 1). The summaries of all of these curations are stamped with the HL GCEP approval date and are live on the ClinGen website (https://search.clinicalgenome.org/kb/gene-validity). The majority of these classifications (105, 64%) were for NSHL, while 59 curations (36%) were for syndromic conditions (Figure 2B). We curated 19 genes with respect to more than one disease and/or inheritance pattern (Supplementary Table 1). Detailed clinical information on 44 syndromic genes where hearing loss is not the presenting feature can be found in Supplementary Table 2.

**Figure 2:**
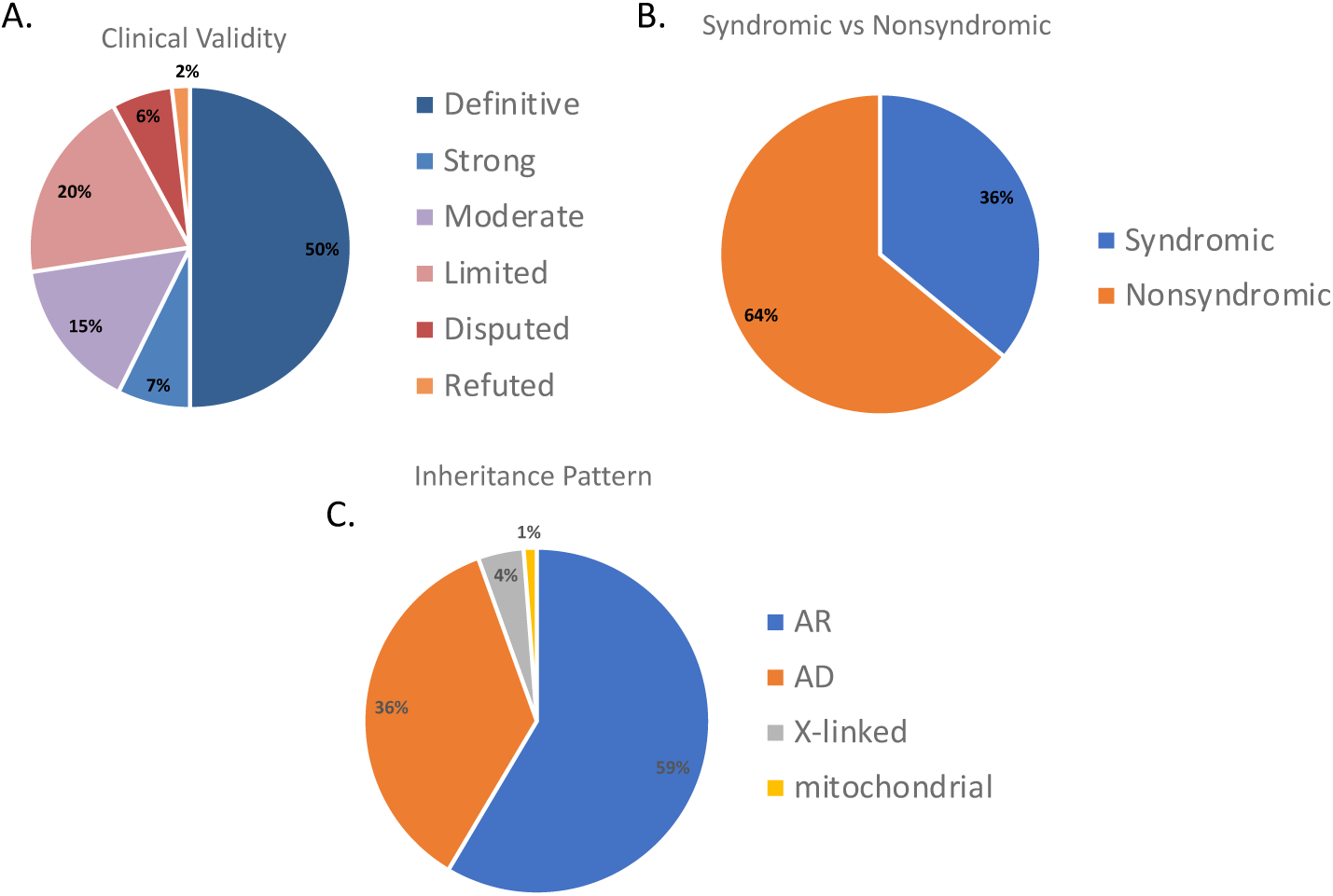
A. The clinical validity of 164 gene-disease pairs; Definitive= 82, Strong=12, Moderate=25, Limited=32, Disputed=10, Refuted= 3. B. Syndromic (N=59) and nonsyndromic (N=105) breakdown of 164 gene-disease pairs. C. Curations split by inheritance pattern; Autosomal Recessive (AR)=96, Autosomal Dominant (AD)=59, X-linked=7, Mitochondrial=2

### Strong and Definitive Gene-Disease Pairs

As per the ClinGen framework,^4^ gene-disease relationships that score 12-18 total points can be Strong or Definitive, the latter category requiring replication over time (>2 publications with convincing evidence over three years after the initial publication). Overall, 82 (50%) gene-disease relationships were Definitive and 12 (7%) were Strong (Figure 2A). Definitive gene-disease pairs were nearly evenly split between syndromic (39/82) and nonsyndromic (43/82). Similarly, Strong gene-disease associations were nearly evenly split with 7/12 syndromic and five nonsyndromic. There were two Strong genes (*CABP2* and *GJB3*) that scored ≥12 points and met the criteria of replication over time, however the experts in the group downgraded them from Definitive to Strong as the aggregate evidence was not convincing enough to be Definitive. *GJB3* was classified as Strong for Erythrokeratodermia variabilis. The expert panel classified an additional four genes (*SLC17A8*, *LARS2*, *MYO3A*, and *CISD2*) as Strong despite only reaching 10.5-11.75 points, based on the total aggregate evidence which was felt to be sufficient to upgrade the classification.

### Moderate Gene-Disease Pairs

We identified 25 (15%) gene-disease pairs with Moderate clinical validity (7-11 points of combined genetic and experimental evidence) (Figure 2A). The Moderate classification typically means that the evidence is promising and more likely to move over time to Strong/Definitive,^9^ but insufficient evidence exists at this time. Of the Moderate gene-disease pairs, 20 were nonsyndromic and five were syndromic. Moderate genes scored 2-8.5 points of genetic evidence and 0.5-6 points of experimental evidence. One gene-disease pair, *MSRB3* and ARNSHL, scored as Strong using the framework point values, however the group determined with expert judgment that the relationship should be downgraded from Strong to Moderate because the only variants identified were homozygous variants in consanguineous families. Another gene, *COL11A2*, was determined to have a Moderate association with both AD and AR NSHL, despite a Definitive association with both AD and AR Otospondylomegaepiphyseal dysplasia (OSMED).

### Limited Gene-Disease Pairs

We identified 32 (20%) gene-disease pairs with Limited clinical validity (0-6 points of combined genetic and experimental evidence) (Figure 2A). Of these, 26 were associated with NSHL and six were associated with syndromic conditions. The Limited genes scored 0.25-8 points of genetic evidence and 0-6 points of experimental evidence. This most often corresponded to an individual proband or a small consanguineous family with a homozygous missense variant. For example, the gene *BDP1* has a Limited relationship with ARNSHL which scored three points. Only one variant was identified, which extends the BDP protein product by 11 amino acids and was found in a homozygous state in a consanguineous family of Qatari descent (NM_018429.2:c.7873T>G (p.Ter2625Glu)). This family had four unaffected individuals and four individuals affected with bilateral, sensorineural, post-lingual onset (ages 2-4 years) progressive hearing loss,^10^ which was scored 0.5 variant points and 2 segregation points. The experimental evidence demonstrates that *BDP1* is expressed in murine endothelial cells of stria vascularis capillaries, and mesenchyme-derived cells and surrounding extracellular matrix around the cochlear duct including the spiral ligament and basilar membrane,^10^ which was scored 0.5 points. While expression evidence suggests that the gene may have cochlear function, it does not prove it is required for function and the segregation evidence does not uniquely implicate this gene given the large linkage interval. Therefore, with only a single family reported, the gene-disease pair resulted in a Limited association.

### Disputed/Refuted/No Evidence Gene-Disease Pairs

The HL GCEP classified 10 (6%) gene-disease pairs as Disputed (Figure 2A). While evidence for these relationships varied, most often, the small amount of case-level evidence available was not scorable. This differentiates these pairs from genes with “No reported evidence” because a disease claim was made in the literature and case-level information was published. However, the Disputed classification indicates that the expert panel reviewed the evidence and disputed the claim due to insufficient or contradictory evidence. For example, *KCNJ10*, a gene included on 11 panels, has been associated with AR Enlarged vestibular aqueduct (EVA) in two probands from one paper. One proband also carries a missense variant in *SLC26A4* that has been reported in ClinVar as Likely Pathogenic by four clinical testing labs (Partners LMM SCV000060075.5; GeneDx SCV000565574.4; ARUP SCV000605152.1; Counsyl SCV000678181.1).^11^ The second proband, with a homozygous *KCNJ10* variant that is present in high frequency in gnomAD, also carries a splice-site variant in *SLC26A4* that has been reported in ClinVar as Pathogenic by the ClinGen Hearing Loss Expert Panel (SCV000840527.1).^11^ The claim for a digenic inheritance of *SLC26A4* and *KCNJ10* is otherwise weak, therefore the HL GCEP approved *KCNJ10* and AR EVA as Disputed.

The HL GCEP classified 3 (2%) gene-disease pairs as Refuted (Figure 2A). This Refuted classification indicates that the expert panel reviewed the evidence and Refuted the claim due to contradictory evidence significantly outweighing evidence supporting the claim. These three pairs were *GJB6* and ARNSHL, *HARS* and Usher syndrome, and *MYO1A* and ADNSHL. For example, *HARS* was first reported to be associated with AR Usher syndrome in 3 individuals.^12,13^ No convincing segregation or functional information has been reported in order to consider the variants as pathogenic or score the reported cases. For example, the first individual was from the Old Order Amish population and was homozygous for a missense variant in *HARS*, but was homozygous for 80 variants in the linkage interval.^12^ The experimental evidence was limited to a functional study of one of the variants that wasn’t scored. Therefore, this gene-disease pair was Refuted. No gene-disease pairs were classified as No Evidence which was unsurprising given the source of genes was clinically offered panels or publications with reported data.

### GTR Panel and ClinVar analysis

When final approved classifications were plotted against the number of panel tests on which they appeared (Figure 3), 82% (58/73) of Definitive genes appeared on 10 or more panels. However, eight Definitive genes were on five panels or fewer. This may indicate a discrepancy in how often labs update the gene content of their panels. Of these eight Definitive genes, five were associated with syndromic hearing loss (*SLC52A2* and Brown-Vialetto-Van Laere syndrome, *DNMT1* and DNMT methylopathy, *BCS1L* and Bjornstad syndrome, *AIFM1* and Auditory neuropathy spectrum, *CLPP* and Perrault syndrome), suggesting that labs may be less likely to include syndromic genes on comprehensive hearing loss panels. Moderate genes were highly variable in their inclusion on panels. Of the 25 Limited genes, 68% (17/25) were on five panels or fewer. Almost half of the Disputed genes (4/9) were on five panels or fewer and of the three Refuted genes, *GJB6* was on all 17 panels, *MYO1A* was on 10 panels, and *HARS* was on three panels.

**Figure 3:**
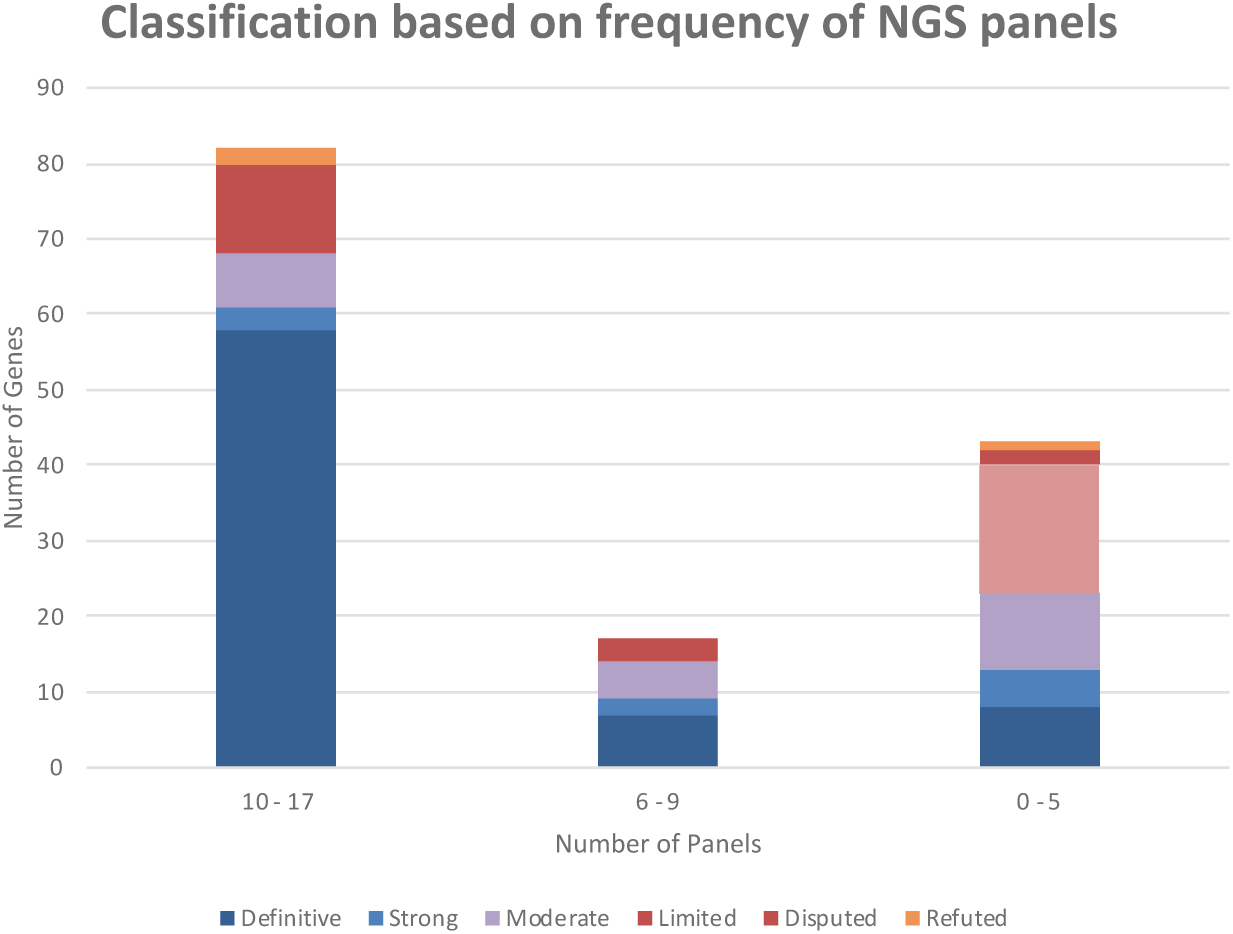
Gene-disease pairs were plotted against the binned number (0-5, 6-9, 10-17) of Next Generation Sequence (NGS) panels on which they appear. These NGS panels were the 17 panels used to assemble a curation gene list per the methods. Genes that were linked with more than one disease were only plotted once with their highest classification. Total gene-disease pairs plotted on this graph N=142

The ClinVar Miner tool^14^ (https://clinvarminer.genetics.utah.edu/) was used to assess the number of variants with “criteria provided” that were submitted to ClinVar with clinical testing as the collection method for each of the Limited, Disputed and Refuted gene-disease pairs (Figure 4). Of the 132 total variants reported in Refuted gene-disease pairs, only one variant was submitted with a clinical significance of Pathogenic. This missense variant in *GJB6* has been submitted to ClinVar as Pathogenic by three clinical testing labs. Two of them submitted it linked with AD Hidrotic ectodermal dysplasia syndrome (Partners LMM SCV000198189.4; GeneDx SCV000321729.6), which has been assessed by the HL GCEP as Definitive gene-disease relationship (Supplementary Table 1). The third clinical testing lab submitted the variant as pathogenic associated with multiple conditions, including AR and AD NSHL (Invitae SCV000767480.1). Therefore, the pathogenic claim cannot be attributed specifically to the GJB6- ADNSHL Refuted gene-disease pair. Of the 116 total variants reported in Limited gene-disease pairs, only two were submitted with a Pathogenic clinical significance. One of these variants was in *KARS* and was scored in the HL GCEP’s Limited curation of *KARS* and ARNSHL (ClinVar Variation ID: 60752). The second variant was in *DCDC2* and was submitted to ClinVar by one lab with two diseases, a syndromic renal condition and nonsyndromic hearing loss condition, and therefore the pathogenic claim cannot be attributed specifically to the DCDC2-ARNSHL Limited gene-disease pair (ClinVar Variation ID: 501347). The majority of variants submitted for Limited 67% (78/116), Disputed 72% (96/134), and Refuted 66% (87/132) gene-disease pairs were of Uncertain clinical significance. Additionally, 29% (34/116) of Limited, 28% (37/134) of Disputed, and 31% (41/132) of Refuted variants were likely benign or benign.

**Figure 4:**
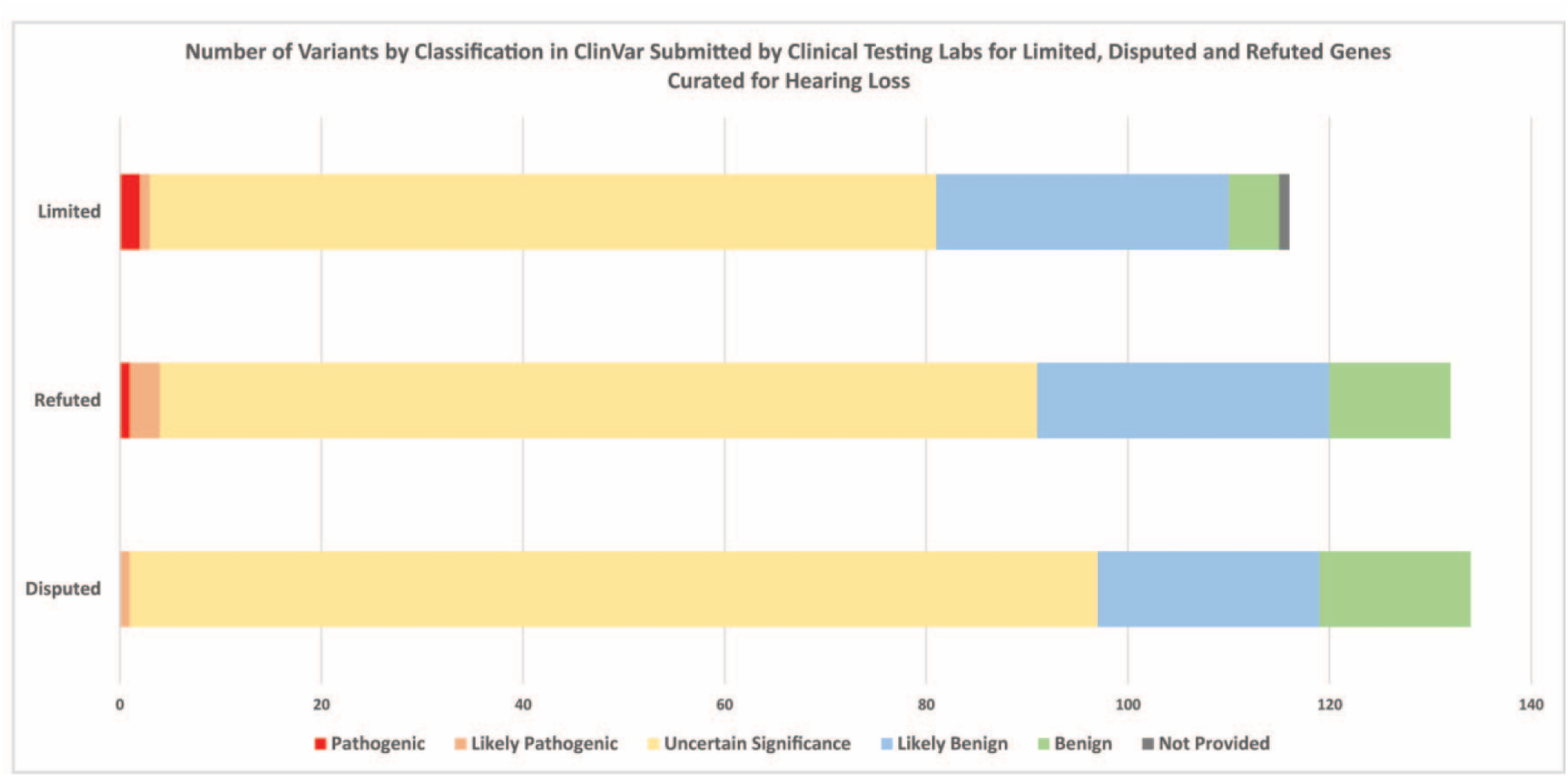
ClinVar miner was used to pull all variants submitted with assertion criteria to ClinVar with the collection method “clinical testing”. If genes had a higher classification in addition to a Limited, Disputed, Refuted classification, only submissions linked to the Limited, Disputed, or Refuted disease entity were counted. Limited (N=25), Refuted (N=3), Disputed (N=10) gene- disease pairs.

## Discussion

We applied the ClinGen clinical validity framework and performed evidence-based curation of 142 genes associated with nonsyndromic and syndromic hearing loss that are included on panels from 17 diagnostic testing laboratories. Several of these genes had more than one disease association that differed by either phenotypic presentation or inheritance pattern, bringing the total number of gene-disease associations that were assessed to 164. The clinical validity classifications for these genes are publicly available: https://search.clinicalgenome.org/kb/gene-validity and listed in Supplementary Table 1.

Of note, roughly a quarter of gene-disease associations (45/164) have a Limited, Disputed or Refuted classification. While Limited associations had scorable human genetic evidence, they often lacked compelling experimental evidence, thus more genetic and/or experimental evidence is needed to meet contemporary criteria for implicating a gene in disease. Furthermore, data have shown that most genes in the Limited category, particularly those that remain Limited for more than five years, do not accumulate evidence in the future to move to a higher classification.^9^ Disputed associations have a disease claim based on human genetic data. However, the evidence for the claim is so minimal that experts dispute it despite not being able to rule out all of the reported evidence. A Refuted classification indicates that there was no scorable genetic evidence supporting the gene-disease claim and **all** prior evidence was refuted (e.g. all reported variants were later found to have high allele frequencies in the general population or later clarified to be in a pseudogene). While the Disputed and Refuted classifications have published claims made using human genetic data, No Evidence gene-disease relationships have no prior claim in the published literature.

One Refuted gene, *GJB6*, appeared on all 17 GTR panels examined, although this was not surprising. Coding variants in *GJB6* are Definitively associated with Clouston syndrome/Hidrotic ectodermal dysplasia, a syndrome characterized by hair loss and skin/nail abnormalities and no hearing loss, but its relationship with ARNSHL has only been documented through large genomic deletions, including *GJB6*-D13S1830 and *GJB6*-D18S1854. These deletions have been identified in *trans* with pathogenic *GJB2* variants in many cases. Specifically, *GJB6*-D13S1830 is a deletion of approximately 309kb of DNA including the 5’ end of *GJB6* and a region upstream of both *GJB6* and the *GJB2* gene, which has been shown to eliminate a *cis*-acting element thereby abolishing expression of the *cis GJB2* allele.^15,16^ Additionally, an independent mouse model with only the coding sequence of *GJB6* deleted and no surrounding sequence deleted had normal hearing, confirming that the regulatory region 5’ of *GJB6*, but not the gene itself, is necessary for normal hearing in mice. Furthermore, many deletions upstream of both *GJB6* and *GJB2* are pathogenic for hearing loss without disruption of *GJB6*.^17,18^ Therefore, the HL GCEP concluded that coding variants in *GJB6* are not associated with hearing loss. The two other Refuted genes, *MYO1A* for NSHL and *HARS* for Usher syndrome were found on 10 and three panel tests, respectively.

Of the 142 genes, 19 were associated with more than one phenotype or inheritance pattern, and the strength of different associations in the same gene varied for several of these genes. For example, four genes (*CDH23*, *MYO7A*, *PCDH15*, *USH1C*) were associated with Usher syndrome type I (USH1) and NSHL (Supplementary Table 1). For *CDH23* and *MYO7A*, associations with both USH1 and NSHL were classified as Definitive, while for *PCDH15* and *USH1C*, the USH1 association was classified as Definitive while NSHL only met a Limited classification. Another example is the *COL11A2* gene which is associated with OSMED and NSHL. Both phenotypes have been associated with recessive and dominant inheritance (Supplementary Table 1), with only the dominant and recessive OSMED relationships meeting criteria for a Definitive classification. Curation of distinct phenotypes and inheritance patterns is important to enable a better prediction of the possible disease presentation and inheritance patterns when novel variants are identified in these genes. However, many of these genes exhibit variable expressivity and age of onset of additional syndromic features which may be missed during initial evaluation, making a determination of the evidence for nonsyndromic associations difficult. In such genes, unless there is a distinct molecular mechanism for syndromic versus nonsyndromic presentations, the syndrome should not be ruled out in patients with a positive genetic result.

As mentioned, diseases associated with one gene were either lumped together or split and curated separately according to the ClinGen Lumping and Splitting Guidelines: (https://www.clinicalgenome.org/site/assets/files/9703/lumping_and_splitting_guidelines_gene_curation_final.pdf). Conditions were lumped if they could not be differentiated by molecular mechanism or inheritance pattern. In many of these cases, the presentations are likely part of a phenotypic spectrum. An example of this is *CHD7* and the relationships with CHARGE syndrome and Kallman syndrome. Because features of CHARGE syndrome have been identified in some Kallman syndrome patients with pathogenic *CHD7* variants, we decided to lump all of these conditions into “CHARGE syndrome” for curation purposes.^19–22^ In addition, the HL GCEP only curated *SLC26A4* for Pendred Syndrome, given the lack of any defining molecular basis for those patients who present without thyroid disease, which could better be considered a phenotype with variable expression.^23,24^ Diseases that were split clearly differed in inheritance pattern, molecular mechanism, or phenotype. For example, *CDH23* was curated for AR Usher syndrome and separately for ARNSHL. These diseases are delineated by variant spectrum. Generally, variants that do not cause full loss of function are associated with NSHL, while loss of function variants are associated with Usher syndrome.^25,26^ Another example of a differentiating molecular mechanism for Usher syndrome and NSHL occurs in *USH1C* where both conditions were also curated separately. Variants in *USH1C* that give rise to Usher syndrome are located in a transcript region expressed in both eye and ear tissue whereas NSHL variants are in regions only expressed in ear tissue.^27^ Therefore, variants that occur in the exons that are present in the eye and ear cause retinitis pigmentosa and hearing loss, while variants only expressed in the ear tissue exclusively cause NSHL.

A major benefit of data sharing beyond classification of variants, is the possibility of strengthening gene-disease relationships. During our curation, three gene-disease associations benefited from the ClinGen community data sharing approach: *OTOG*, *GRHL2* and *ESRRB*. Based on the literature, all three genes had only enough evidence to be classified as Moderate; however, after obtaining case observation evidence from several clinical labs that submitted variants in these genes to ClinVar, the classifications of *OTOG* and *ESRRB* were upgraded to Definitive and *GRHL2* was upgraded to Strong. These examples highlight the importance of ClinVar submission as a mechanism to strengthen both variant and gene level evidence.

In conclusion, the HL GCEP used the ClinGen clinical validity framework to perform evidence-based curation of 142 genes associated with nonsyndromic and syndromic hearing loss, consisting of 164 gene-disease pairs with 82 Definitive (50%), 12 Strong (7%), 25 Moderate (15%), 32 Limited (20%), 10 Disputed (6%), and 3 Refuted (2%) classifications (Figure 2A, Supplementary Table 2). The summaries of all curations are live on the ClinGen website (https://search.clinicalgenome.org/kb/gene-validity). ACMG is currently developing standards for diagnostic gene panel development. We suggest to be consistent with those guidelines and include only those genes with at least a Moderate level of evidence in diagnostic tests for hearing loss. Furthermore, we recommend inclusion of at least the syndromic genes listed in Supplementary Table 1 given the possibility of missing the syndromic diagnosis due to delayed onset, variable expressivity or subtle presentations of non-hearing loss features. This approach will serve to optimize the clinical sensitivity of testing while reducing the rate of VUSs due to genes with insufficient evidence.

## Acknowledgements

Research reported in this publication was supported by the National Human Genome Research Institute (NHGRI) under award number U41HG006834. The content is solely the responsibility of the authors and does not necessarily represent the official views of the National Institutes of Health.

## Supplementary Figure Legend

Supplementary Figure 1: ClinVar miner was used to pull all variants submitted with assertion criteria to ClinVar with the collection method “clinical testing”. If genes had a higher classification in addition to a Limited, Disputed, Refuted classification, only submissions linked to the Limited, Disputed, or Refuted disease entity were counted. A. List of Limited genes (N=25) and classifications. B. List of Disputed (N=10) and Refuted (N=3) genes.

