## Supplementary Figure 1 for "ClinGen Expert Clinical Validity Curation of 164 Hearing Loss Gene-Disease Pairs"

### MT-RNR1- mitochondrial non-syndromic sensorineural deafness with susceptibility to aminoglycoside exposure- *Mitochondrial inheritance*

**Classification Owner:** Hearing Loss EP
**Calculated classification:** Definitive
**Modified classification:** Definitive
**Reason for modified classification:** None
SOP: Gene Clinical Validity Standard Operating Procedures (SOP), Version 6

**Classification status:**
**Date classification saved:** 2018 Oct. 5, 11:37 am
**Replication Over Time:** Yes
**Contradictory Evidence?** Proband: **No**, Experimental: **No**
Disease: mitochondrial non-syndromic sensorineural deafness with susceptibility to aminoglycoside exposure

### Evidence Summary

The MTRNR1 gene has been associated with maternally-inherited, aminoglycoside-induced hearing loss using the ClinGen Clinical Validity Framework as of 7/16/2018. MTRNR1 variants have repeatedly demonstrated that they cause a predisposition to developing hearing loss that can onset or worsen with aminoglycoside exposure. Hearing loss in the absence of aminoglycoside exposure has also been reported but was not assessed separately from this curation. This association was made using case-level data, case-control data and experimental evidence. At least 9 unique variants (1555A>G, 1494C>T, 827A>G, 1095T>C, 961delT, 961delTinsC2_7, 961T>G, 961T>C, 961insC) have been reported in humans with hearing loss, though the alterations at position 1095, 961, and 827 have also been seen in high frequencies in the general population or in haplogroups in the GenBank database (www.mitomap.org, Barbarino et al. 2016). Most of the cases with these variants were identified as homoplasmic though some heteroplasmic cases have been identified (Thyagarajan 2000, Tessa 2001). MTRNR1 was first associated with this disease in humans as early as 1993 (Prezant et al.). The association is seen in more than 500 individuals with aminoglycoside exposure in 60 publications as of 2016 (Barbarino 27654872). Variants in this gene segregated with disease in more than 20 additional family members (Zhao 2004, Mutai 2017). The 1555A>G variant has been identified in both homoplasmic and heteroplasmic occurances and heteroplasmic load levels have been correlated to the severity of the hearing loss within families (Ballana 2008, del Castillo 2003). Of note, some reports suggest that variants such as the established 1555A>G variant can cause hearing loss without aminoglycoside exposure, but these associations are not as clear due to the self-reported nature of aminoglycoside usage data and incomplete penetrance in these families (Li 2005, Bykhovskaya 1998). The mechanism for disease is thought to be that alterations in conserved regions disrupt the secondary structure of the mitochondrial 12S rRNA leading to altered mitochondrial function that leads to susceptibility of the ototoxicity of aminoglycosides (Zhao 2004). This gene-disease association is supported by in vitro assays showing lymphoblastoid patient mitochondria have altered translational fidelity and oxygen consumption (Zhao 2005, Zhao 2005, Hobbie 2008, Guan 1996). Per criteria outlined by the ClinGen Lumping and Splitting Working Group, we found no difference in molecular mechanism underlying the disease entities: (1) nonsyndromic hearing loss; (2) aminoglycoside induced hearing loss. In summary, MTRNR1 is definitively associated with maternally-inherited, aminoglycoside-induced hearing loss. This has been repeatedly demonstrated in both the research and clinical diagnostic settings, and has been upheld over time. This classification was approved by the Hearing Loss Gene Curation Working Group on 7/17/2018.

### Calculated Classification Matrix

| Evidence Type | | | | | | Count | Total Points | Points Counted |
|---|---|---|---|---|---|---|---|---|
| Genetic Evidence | Case-Level | Variant | Autosomal Dominant OR X-linked Disorder | Proband with other variant type with some evidence of gene impact | | 7 | 2.6 | 2.6 |
| | | | | Proband with predicted or proven null variant | | 0 | 0 | 0 |
| | | | | Variant is *de novo* | | 2 | 3.5 | 3.5 |
| | | | Autosomal Recessive Disorder | Two variants (not predicted/proven null) with some evidence of gene impact in *trans* | | 0 | 0 | 0 |
| | | | | Two variants in *trans* and at least one *de novo* or a predicted/proven null variant | | 0 | 0 | 0 |
| | | Segregation | | | **Summed LOD** | **Family Count** | 3 | 3 |
| | | | | Candidate gene sequencing | 0 | 0 | | |
| | | | | Exome/genome or all genes sequenced in linkage region | 5 | 1 | | |
| | | | | **Total Summed LOD Score** | 5 | | | |
| | **Case-Control** | | | | | 1 | 3 | 3 |
| | **Genetic Evidence Total:** | | | | | | | **12.1** |
| Experimental Evidence | **Functional** | | | Biochemical Functions | | | 0 | 0 |
| | | | | Protein Interactions | | | 0 | |
| | | | | Expression | | | 0 | |
| | **Functional Alternation** | | | Patient Cells | | 4 | 2 | 2 |
| | | | | Non-patient cells | | | 0 | |
| | **Models** | | | Non-human model organism | | | 0 | 0 |
| | | | | Cell culture model | | | 0 | |
| | **Rescue** | | | Rescue in human | | | 0 | 0 |
| | | | | Rescue in non-human model organism | | | 0 | |
| | | | | Rescue in cell culture model | | | 0 | |
| | | | | Rescue in patient cells | | | 0 | |
| | **Experimental Evidence Total:** | | | | | | | **2** |
| **Total Points** | | | | | | | | **14.1** |

**Genetic Evidence: Case Level (variants, segregation)**

| Label | Variant type | Variant | Reference | Proband sex | Proband age | Proband ethnicity | Proband phenotypes | Segregations | | | | | Proband previous testing | Proband methods of detection | Score status | Proband points (default points) | Reason for changed score |
|---|---|---|---|---|---|---|---|---|---|---|---|---|---|---|---|---|---|
| | | | | | | | | # Aff | # Unaff | LOD score | Counted | Sequencing | | | | | |
| Mutai_1 | Proband with other variant type with some evidence of gene impact | m.1555A>G | Mutai H, et al., 2017, PMID: 28320335 | Male | | | **HPO term(s):** Severe sensorineural hearing impairment Progressive hearing impairment **free text:** steeply sloping progressive HL | 4 | – | – | – | | "Mutai et al evaluated for bilateral and symmetric SNHL ≥ 4 family members with SNHL with a maternal trait of inheritance in ≥ 2 generations, 3) onset of SNHL before the age of 40 years, 4) high-frequency SNHL, and 5) no record of environmental factors related to SNHL." | | Score | **0.5 pts** (variant) **3 points** (segregation) | see screenshots of mito databases below, generally absent/rare frequency. Due to the unique inheritance pattern, it appears that this is definitely the cause of the hearing loss for all these individuals (aminoglycoside exposure not noted) |
| Mutai_2 | Proband with heteroplasmy trend | m.1555A>G | Mutai H, et al., 2017, PMID: 28320335 | Female | 21 | | **HPO term(s):** Severe sensorineural hearing impairment Progressive hearing impairment **free text:** steeply sloping progressive HL | – | – | – | – | | "Mutai et al evaluated for bilateral and symmetric SNHL ≥ 4 family members with SNHL with a maternal trait of inheritance in ≥ 2 generations, 3) onset of SNHL before the age of 40 years, 4) high-frequency SNHL, and 5) no record of environmental factors related to SNHL." | | Score | **2 pts** | Each of these individuals had at least 4 family members with SNHL and maternal trait of inheritance for at least 2 generations but seqeuncing of the other family members was not conducted. see screenshots of mito databases below, generally low frequency. Due to the unique inheritance pattern, it appears that this is definitely the cause of the hearing loss for all these individuals |
| Mutai 4 | Proband with heteroplasmy trend | m.1555A>G | Mutai H, et al., 2017, PMID: 28320335 | Female | | | | – | – | – | – | | "Mutai et al evaluated for bilateral and symmetric SNHL ≥ 4 family members with SNHL with a maternal trait of inheritance in ≥ 2 generations, 3) onset of SNHL before the age of 40 years, 4) high-frequency SNHL, and 5) no record of environmental factors related to SNHL." | | Score | **1.5 pts** | see screenshots of mito databases below, generally low frequency. Only 3 of these indiviudals met all 5 criteria but they still all carried the variant and had HL. This individual did not meet al the criteria |

| | | | | | | | | | | | | | | | | | |
|---|---|---|---|---|---|---|---|---|---|---|---|---|---|---|---|---|---|
| Chai_4 | Proband with heteroplasmy trend | m.1555A>G | Chai Y, et al., 2014, PMID: 25251670 | Unknown | | | **free text:** bilateral permanent non-dominant hearing loss | _ | _ | _ | _ | | Dominant inheritance excluded, had confirmed maternal inheritance pattern | | Score | **1.5 pts** | see screenshots of mito databases below, generally low frequency. 5 individuals (4.4% of the 114 tested probands with hearing loss had the MTRNR1 variation. It is an extremely common cause of HL. |
| 3 cases with m.827A>G | Proband with other variant without evidence for impact | m.827A>G | Li Z, et al., 2005, PMID: 15841390 | Unknown | | | **free text:** aminoglycoside induced hearing loss | _ | _ | _ | _ | | | **Method 1:** PCR **Description of genotyping method**: PCR sequencing of MT-RNR1 | Not scored | **0 pts** | High conservation in species, found in multiple individuals with AG-induced HL but found in 100% frequency in several South East Asian/ Native American haplogroups in mitomap. 28% in Arabia/Horn of Africa haplogroup |
| Li case with m.1005T>C | Proband with other variant without evidence for impact | m.1005T>C | Li Z, et al., 2005, PMID: 15841390 | Unknown | | | **free text:** aminoglycoside induced hearing loss | _ | _ | _ | _ | | | | Not scored | **0 pts** | Found in 100% of several haplogroups incuding F2a, A11a (Asia, Tibetan plateau) cannot be scored |
| Li Case w/ m.1116A>G | Proband with other variant without evidence for impact | m.1116A>G | Li Z, et al., 2005, PMID: 15841390 | Unknown | | | | _ | _ | _ | _ | | | | Downgraded | **0.1 pts** (variant) | High conservation in other species and highest frequency in HG branch was 0.75% Can't score |

| | | | | | | | | | | | | | | | | | |
|---|---|---|---|---|---|---|---|---|---|---|---|---|---|---|---|---|---|
| Zhao family proband IV:21 | Proband with other variant type with some evidence of gene impact | m.1494C>T | Zhao H, et al., 2004, PMID: 14681830 | Male | | | **HPO term(s):** Profound sensorineural hearing impairment Childhood onset sensorineural hearing impairment **free text:** Proband has a profound sensorineural hearing impairment that appears to have been induced by aminoglycosides at 1.5 years | 20 | – | **Calculated:** 5 | Yes | Exome/genome or all genes sequenced in linkage region | This paper conducted detailed phenotyping of all affected members of the family and noted aminoglycoside usage for each indiviudal. It appears that aminoglycosides worsen the hearing impairment. | **Description of genotyping method:** mitochondrial genome sequencing | Score | **0.5 pts** (variant) | SCORE VARIANT AS 0.5 PTS, "Examined known mtDNA mutations associated with deafness by PCR and then conducted PCR w/ 24 fragments spanning the entire mitochondrial genome of the proband. Did not detect the A1555G, mutation in the 12S rRNA gene or the A7445G, T7510C, and T7511C mutations in the t- RNASer(UCN) gene in those patients. Given that this is essentially sequencing every gene in a candidate region, this was eligable for the full 3 points of segregation. Additionally, functional evidence supports the impact of the variant on the protein." |
| Wang 4 generation chinese family. Proband II-2 | Proband with other variant type with some evidence of gene impact | m.1494C>T | Wang Q, et al., 2006, PMID: 16380089 | Female | **Age of Onset:** 6 Years | | **HPO term(s):** Profound sensorineural hearing impairment Persistent stapedial artery **free text:** Administered with streptomycin for fever at the age of 6 years and began suffering hearing loss one month after drug admin. | 3 | – | – | – | | Family members who were administered aminoglycosided had hearing loss, and some who were not administered had HL but this does not rule out exposure. Amplified DNA from the 12S rRNA and tRNASerUCN genes. DNA from 4 matrilineal relatives, proband II-2, affected daughter III-2, affected male II-3 and unaffected male II-5 as well as two unrelated Chinese controls. | | Score | **0 pts** | Same variant as above, this paper did not sequence enough individuals for segregation information to be counted however the variant can still be scored. |
| | | | | | | | | | | | | | | | | **Total Points:** | **9.10** |

**Genetic Evidence: Case-Control**

| Label | Reference (PMID) | Disease (Case) | Study Type | Detection method (Case) | Power: # of cases genotyped/ sequenced | Power: # of controls genotyped/ sequenced | Bias confounding | Statistics: Cases with variant in gene/ all cases genotyped/sequenced | Statistics: Controls with variant in gene/ all cases genotyped/sequenced | Statistics: Test statistic: value | Statistics: p-value | Statistics: Confidence interval | Points |
|---|---|---|---|---|---|---|---|---|---|---|---|---|---|
| 813 Probands from Hubei province | Chen G, et al., 2011, PMID: 21777984 | mitochondrial non-syndromic sensorineural deafness with susceptibility to aminoglycoside exposure (MONDO: 0010799) | Single variant analysis | | 813 | 123 | | 96/813 | 0/123 | **Other:** Chi Squared with yates correction - 266.151 | 0.00001 | | 3 |
| | | | | | | | | | | | | **Total points:** | **3.00** |

| Experimental Evidence | | | | | | |
|---|---|---|---|---|---|---|
| **Label** | **Experimental category** | **Reference** | **Explanation** | **Score status** | **Points (default points)** | **Reason for changed score** |
| Lymphoblastoid cell lines | **Functional Alteration:** Patient cells | Zhao H, et al., 2004, PMID: 14681830 | Lymphoblastoid cell lines from 6 individuals with the m.1494C>T variant grown in a medium in the presence of aminoglycoside drugs and in their absence and found that there was an effect on cell growth as well as oxygen consumption rate with sensitivity to aminoglycoside paromomycin | Score | **0.5** (1) | Slightly downgraded the points from the 1 default given to patient cell lines, due to lack of direct implication in HL, but still felt that this was indicative of the trend of aminoglycoside ototoxicity seen in humans and gave 0.5 pts |
| Patient lymphoblastoid cell lines | **Functional Alteration:** Patient cells | Guan MX, et al., 1996, PMID: 8817331 | Lymphoblastoid cell lines sensitive to aminoglycoside exposure cultivated from 9 individuals with 1555A>G variant from an Arab-Isreaeli family and 13 individuals without the variant. Consistent with disease presentation | Score | **0.5** (1) | Lymphoblastoid cell lines sensitive to aminoglycoside exposure cultivated from 9 individuals with 1555A>G variant from an Arab-Isreaeli family and 13 individuals without the variant. Consistent with disease presentation |
| Patient cell lines 1494C>T | **Functional Alteration:** Patient cells | Zhao H, et al., 2004, PMID: 14681830 | 9 patient cell lines with the 1494C>T variant showed that they had significantly less labeled translation products than the control cells indicating that the variants in the mitochondrial gene have a wide array of issues impacting mitochondrial function. They also found thta there was a variable decrease in the O2 consumption rate, that was also significant. Sensitivity to aminoglycosides was measured and confirmed | Score | **0.5** (1) | patient cell lines with lack of direct implication in HL |
| Mutant strains of M.smegmatis | **Functional Alteration:** Non-patient cells | Hobbie SN, et al., 2008, PMID: 18308926 | Constructed mutant strains of M.smegmatis and isolated the ribosomes to show that variants cause an alteration that compromises translational fidelity of ribosome. This phenomenon was significantly increased w/ aminoglycosides | Score | **0.5** (.5) | |
| | | | | **Total points:** | **2.00** | |
