## Supplementary Figure 2 for "ClinGen Expert Clinical Validity Curation of 164 Hearing Loss Gene-Disease Pairs"

### MT-TS1- mitochondrial non-syndromic sensorineural deafness with susceptibility to aminoglycoside exposure- *Mitochondrial inheritance*

**Classification Owner:** Hearing Loss EP
**Calculated classification:** Definitive
**Modified classification:** None
**Reason for modified classification:** None
**SOP:** Gene Clinical Validity Standard Operating Procedures (SOP), Version 6

**Classification status:**
**Date classification saved:** 2018 Dec 20, 8:36 am
**Replication Over Time:** Yes
**Contradictory Evidence?** Proband: **No**, Experimental: **No**
Disease: mitochondrial non-syndromic sensorineural deafness with susceptibility to aminoglycoside exposure

### Evidence Summary

MT-TS1 was first reported in relation to mitochondrial nonsyndromic hearing loss in 1993 (Prezant et al., 7689389). At least seven variants have been reported in humans (7445A>G, 7445A>C, 7445A>T, 7472insC, 7505T>C, 7510T>C, 7511T>C). Evidence supporting this gene-disease relationship includes case-level data, segregation data, and experimental data. Association is seen in at least 12 probands in 11 publications (PMIDs: 12461693, 8572257, 10094190, 10371545, 20153673, 27530448, 28320335, 15292920, 12471220, 18639500, 7581383). Variants in this gene segregated with disease in 92 additional family members. The gene-disease association is supported by in vitro functional assays (PMIDs: 18398437, 15694374, 15336535). Variants in this gene have been implicated in additional phenotypes including palmoplantar keratoderma with deafness, mitochondrial cytochrome c oxidase deficiency, exercise intolerance, MERRF/MELAS overlap syndrome and ptosis, hypotonia, seizures, and dilated cardiomyopathy. These are outside the scope of this assessment. In summary, MT-TS1 is definitively associated with mitochondrial nonsyndromic hearing loss. This has been repeatedly demonstrated in both the research and clinical diagnostic settings, and has been upheld over time. This classification was approved by the ClinGen Hearing Loss Working Group on 7/17/2018.

### Calculated Classification Matrix

| Evidence Type | | | | | Summed LOD | Count | Total Points | Points Counted |
|---|---|---|---|---|---|---|---|---|
| Genetic Evidence | Case-Level | Variant | Autosomal Dominant OR X-linked Disorder | Proband with other variant type with some evidence of gene impact | | 12 | 6 | 6 |
| | | | | Proband with predicted or proven null variant | | 0 | 0 | 0 |
| | | | | Variant is *de novo* | | 0 | 0 | 0 |
| | | | Autosomal Recessive Disorder | Two variants (not predicted/proven null) with some evidence of gene | | 0 | 0 | 0 |
| | | | | Two variants in *trans* and at least one *de novo* or a predicted/proven | | 0 | 0 | 0 |
| | | Segregation | | | **Summed LOD** | **Family Count** | 3 | 3 |
| | | | | Candidate gene sequencing | | | | |
| | | | | Exome/genome or all genes sequenced in linkage region | 7.83 | 1 | | |
| | | | | **Total Summed LOD Score** | | | | |
| | **Case-Control** | | | | | | | |
| | **Genetic Evidence Total:** | | | | | | | **9** |
| Experimental Evidence | Functional | | | Biochemical Functions | | 0 | 0 | 0.5 |
| | | | | Protein Interactions | | 0 | 0 | |
| | | | | Expression | | 0.5 | 0 | |
| | Functional Alternation | | | Patient Cells | | 0 | 0 | 1.5 |
| | | | | Non-patient cells | | 3 | 1.5 | |
| | Models | | | Non-human model organism | | 0 | 0 | 0 |
| | | | | Cell culture model | | 0 | 0 | |
| | Rescue | | | Rescue in human | | 0 | 0 | |
| | | | | Rescue in non-human model organism | | 0 | 0 | |
| | | | | Rescue in cell culture model | | 0 | 0 | |
| | | | | Rescue in patient cells | | 0 | 0 | |
| | **Experimental Evidence Total:** | | | | | | | **2** |
| **Total Points** | | | | | | | | **11** |

| Genetic Evidence: Case Level (variants, segregation) | | | | | | | | | | | | | | | | | |
|---|---|---|---|---|---|---|---|---|---|---|---|---|---|---|---|---|---|
| Label | Variant type | Variant | Reference | Proband sex | Proband age | Proband ethnicity | Proband phenotypes | Segregations | | | | | Proband previous testing | Proband methods of detection | Score Status | Proband points (default points) | Reason for changed score |
| | | | | | | | | # Aff | # Unaff | LOD score | Counted | Sequencing | | | | | |
| Family 1 Proband | Proband with other variant type with some evidence of gene impact | m.7511T>C | Chapiro E, et al., 2002, PMID: 12461693 | Female | **Age of Report:** 23 Years | | **HPO term(s):** Severe sensorineural hearing impairment | 5 | 3 | **Calculated:** 1.8 | No | | | **Method 1:** PCR | 0.5 | | Variant is absent from mtDB and MitoTIP in silico tool predicts likely pathogenic. |
| F-G Family 1 Proband | Proband with other variant type with some evidence of gene impact | m.7445A>G | Fischel-Ghodsian N, et al., 1995, PMID: 8572257 | Female | **Age of Report:** 23 Years | | **HPO term(s):** Progressive sensorineural hearing impairment Severe sensorineural hearing impairment Adult onset sensorineural hearing impairment | 8 | 2 | _ | _ | | | **Method 1:** PCR; **Method 2:** Sanger sequencing | 0.5 | | Variant is absent from mtDB. In HmtDB, variant is absent from 'Normal' individuals and present in 0.03% 'Patient individuals. MitoTIP in silico tool does not have a prediction. Family is primarily homosplasmic. |
| Verhoeven Family 1 Proband | Proband with other variant type with some evidence of gene impact | m.7471_7472insC | Verhoeven K, et al., 1999, PMID: 10094190 | Male | **Age of Report:** 68 Years | | **HPO term(s):** Progressive sensorineural hearing impairment | 27 | 3 | **Calculated:** 7.83 | Yes | | | **Method 1:** PCR | 0.5 | | Variant is absent from mtDB and HmtDB. MitoTIP in silico tool does not have a prediction. Family is primarily homosplasmic. |
| Sue/Friedman Family Proband | Proband with other variant type with some evidence of gene impact | m.7511T>C | Sue CM, et al., 1999, PMID: 10371545 | Female | | Not Hispanic or Latino | **HPO term(s):** Bilateral sensorineural hearing impairment Progressive sensorineural hearing impairment **free text**: variable onset HL, COX deficiency in muscles | 16 | 4 | _ | _ | | 10340654; Friendman et al. 1999 | **Method 1:** PCR | 0.5 | | Variant is absent from mtDB and MitoTIP in silico tool predicts likely pathogenic. |

| | | | | | | | | | | | | | | | | | |
|---|---|---|---|---|---|---|---|---|---|---|---|---|---|---|---|---|---|
| Hutchin Family 1 Proband | Proband with other variant type with some evidence of gene impact | m.7510T>C | Hutchin TP, et al., 2000, PMID: 10978361 | Male | **Age of Diagnosis:** 15 Months | Not Hispanic or Latino | **HPO term(s):** Profound sensorineural hearing impairment **free text:** asymmetric hearing loss | 3 | – | – | – | | | **Method 1:** PCR | 0 | | not scored because of unspecified/heteroplasmy |
| Tang Family Proband | Proband with other variant type with some evidence of gene impact | NC_012920.1:m.7505T>C | Tang X, et al., 2010, PMID: 20153673 | Male | **Age of Onset:** 3 Years | Not Hispanic or Latino | **HPO term(s):** Bilateral sensorineural hearing impairment Severe sensorineural hearing impairment | 7 | 2 | **Calculated:** 1.8 | No | | | **Method 1:** Genotyping; **Method 2:** Sanger sequencing | 0.5 | | Variant is absent from mtDB and HmtDB. MitoTIP in silico tool predicts possibly pathogenic. |
| TCR26 | Proband with other variant type with some evidence of gene impact | m.7444G>A | Subathra M, et al., 2016, PMID: 27530448 | Female | | Not Hispanic or Latino | **HPO term(s):** Severe sensorineural hearing impairment Prelingual sensorineural hearing impairment Global developmental delay | – | – | – | – | | | **Method 1**: Sanger sequencing | 0 | | not scored because this variant is present in a high number of control individuals. |
| AJ7 | Proband with other variant type with some evidence of gene impact | m.7471_7472insC | Subathra M, et al., 2016, PMID: 27530448 | Female | **Age of Report:** 28 Years | | **HPO term(s):** Profound sensorineural hearing impairment Prelingual sensorineural hearing impairment | – | – | – | – | | | **Method 1**: Sanger sequencing | 0.5 | | Variant is absent from mtDB and HmtDB. MitoTIP in silico tool does not have a prediction. Family is primarily homosplasmic. |
| BJ303-III-1 | Proband with other variant type with some evidence of gene impact | m.7445A>T | Chen J, et al., 2008, PMID: 18639500 | Female | **Age of Onset:** 1 Years | | **HPO terms(s):** Severe sensorineural hearing impairment | – | – | – | – | | | **Method 1:** Sanger sequencing | 0.5 | | Variant is absent from mtDB and HmtDB. Found in 3 of 65 (3.08%) F4b haplogroup in Mitobank. |
| Mutai Family Proband | Proband with other variant type with some evidence of gene impact | m.7511T>C | Mutai H, et al., 2017, PMID: 28320335 | Unknown | | | **HPO term(s):** Moderate sensorineural hearing impairment | 5 | 8 | **Calculated:** 1.2 | No | | | **Method 1:** Sanger sequencing **Description of genotyping method:** mtDNA analysis on 2 rRNA and 22 tRNA genes | 0.5 | | Variant is absent from mtDB and MitoTIP in silico tool predicts likely pathogenic. |

| | | | | | | | | | | | | | | | | | |
|---|---|---|---|---|---|---|---|---|---|---|---|---|---|---|---|---|---|
| Jacobs proband | Proband with other variant type with some evidence of gene impact | m.7445A>G | Jacobs HT, et al., 2005, PMID: 15292920 | Unknown | | | | _ | _ | _ | _ | | | **Method 1:** PCR | 0.5 | | Variant is absent from mtDB. In HmtDB, variant is absent from 'Normal' individuals and present in 0.03% 'Patient individuals. MitoTIP in silico tool does not have a prediction. |
| BJ304-III-4 | Proband with other variant type with some evidence of gene impact | m.7445A>C | Chen J, et al., 2008, PMID: 18639500 | Male | **Age of Report**: 19 Years | | **HPO term(s):** Profound sensorineural hearing impairment | _ | _ | _ | _ | | | **Method 1**: Sanger sequencing | 0.5 | | Variant is absent from mtDB. In HmtDB, variant is present in 0.06% of 'Normal' individuals and 0% in 'Patient" individuals. MitoTIP in silico tool does not have a prediction. |
| de Castillo Proband | Proband with other variant type with some evidence of gene impact | m.7510T>C | del Castillo FJ, et al., 2002, PMID: 12471220 | Female | | | **HPO term(s):** Postlingual sensorineural hearing impairment Moderate sensorineural hearing impairment Moderate sensorineural hearing impairment | 21 | _ | _ | _ | | | **Description of genotyping method**: 148 Spanish families with maternal NSHL inheritence pattern tested for m.1555A>G, m.1095T>C and m.7510T>C variants. | 0.5 | | The variant is absent from mtDB and HmtDB. MitoTIP on silico tool predicts possibly pathogenic. The variant is observed in 21 individuals with maternal pattern inheritance of nonsyndromic hearing loss. |
| Tiranti Proband | Proband with other variant type with some evidence of gene impact | m.7471_7472insC | Tiranti V, et al., 1995, PMID: 7581383 | Male | **Age of Onset**: 25 Years | | **HPO term(s):** Progressive sensorineural hearing impairment Tinnitus Gait ataxia Dysarthria Generalized hypotonia Reduced tendon reflexes **free text:** hypodiadochokinesia | 8 | _ | _ | _ | | | | 0.5 | | Variant is absent from mtDB and HmtDB. MitoTIP in silico tool does not have a prediction. |
| | | | | | | | | | | | | | | **Total Points:** | 6 | | |

| Genetic Evidence: Case Level (family segregation information without proband data or scored proband data) |
|---|
| No segregation evidence for a Family without a proband was found. |

| Genetic Evidence: Case-Control |
|---|
| No scored Case-Control evidence was found |

**Experimental Evidence**

| Label | Experimental category | Reference | Explanation | Score status | Points (default points) | Reason for changed score |
|---|---|---|---|---|---|---|
| Swalwell Expression | **Expression** B | Swalwell H, et al., 2008, PMID: 18398437 | As the level of m.7472insC increases, the steady-state level of mt-tRNA^ser(UCN) decreases. Cells harboring >50% m.7472insC variant exhibit barely detectable transcript levels. Suggests that both m.7472insC and m.7473A>C mutations act synergistically in vitro, with the suppressive effect of the m.7472A>C variant regulated at the level of transcription by the m.7472insC variant amount. | Score | **0.5** (0.5) | |
| Swalwell Non-Patient Cells | **Functional Alteration:** Non-patient cells | Swalwell H, et al., 2008, PMID: 18398437 | clones with the m.7472A>C variant alone had marked respiratory deficiency, whereas clones with both mutations respired at rates comparable to controls | Score | **0.5** (0.5) | |
| Li Non-Patient Cells | **Functional Alteration:** Non-patient cells | Li X, et al., 2005, PMID: 15694374 | The amount of tRNA(Ser(UCN)) in mutant cells decreased compared to control cells. Aminoacylation capability test show a 22% reduction in efficiency of charging in mutated tRNA(Ser(UCN)) compared to controls. | Score | **0.5** (0.5) | |
| Toompuu Non-Patient Cells | **Functional Alteration:** Non-patient cells | Toompuu M, et al., 2004, PMID: 15336535 | extracted and deacetylated RNA from the cell cybrids and found over 11% of all tRNA(Ser(UCN)) molecules from 7472insC cells had incorrect 5' and/or 3' termini. Most commonly, one extra 5' terminal nucleotide (U) added. Errors on the 3' side were more diverse. Authors further found that the incorrect 5' and 3' termini lead to improper tRNA processing | Score | **0.5** (0.5) | |
| | | | | **Total points:** | **2.00** | |
