## Supplementary Table 3 for "ClinGen Expert Clinical Validity Curation of 164 Hearing Loss Gene-Disease Pairs"

Supplementary Table 3: HL GCEP Scoring Specifications

Genetic Evidence

| **Scenario** | **HL GCEP Modified Pts.** |
| --- | --- |
| Homozygous missense variants in consanguineous families with evidence of pathogenicity | 0.25-0.5 |
| Homozygous, non-consanguineous LOF variants | 1-2 |
| Non-truncating/null variants with minimal or no evidence of pathogenicity | 0-0.1 |
| Truncating variants predicted to **not** undergo nonsense mediated decay | 0.5-1.5 |
| *De novo* variants in cases when the hearing loss severity was mild/undisclosed/unspecific due to possible familial inheritance/heterogeneity of the disorder | 0-1 |
| Small-indels with the required functional/segregation/case data to implicate pathogenicity | 0.5-1 |
| Variants reported multiple times within the same ethnic group or country of origin were only scored fully once according to the available evidence. | Full |
| - Subsequent probands were downgraded by at least half of the default score if there was evidence of the probands being unrelated. | Half of default |
| - Subsequent probands were not scored if there was no evidence that the probands were unrelated. | 0 |
| Experimental findings that are used to provide evidence for the scoring of a variant were not scored again under the category of Experimental evidence, so as not to overinflate the scoring of those pieces of evidence. | |

Experimental Evidence

| **Scenario** | **HL GCEP Modified Pts.** |
| --- | --- |
| Convincing mouse model that is consistent in phenotype and mode of inheritance to what has been observed in human patients | 2 |
| Zebrafish, yeast and Model Systems that are less informative for hearing loss phenotypes | 1-1.5 max |
| Expression evidence that is replicated by multiple authors was only scored once. Subsequent expression studies that introduce new, substantive evidence were scored. | 0.5 |
| Syndromic mouse models that were scored for syndromic hearing loss were not scored for the split out nonsyndromic hearing loss curations | |

Abbreviations: HL GCEP, hearing loss gene curation expert panel

LOF, loss of function

Pts., points
